# C9ORF72 deficiency results in degeneration of the zebrafish retina *in vivo*

**DOI:** 10.1101/2023.10.19.563041

**Authors:** Natalia Jaroszynska, Andrea Salzinger, Themistoklis M. Tsarouchas, Catherina G. Becker, Thomas Becker, David A. Lyons, Ryan B. MacDonald, Marcus Keatinge

## Abstract

G4C2 Hexanucleotide repeat expansions within the gene *C9ORF72* are the most common cause of the neurodegenerative diseases Amyotrophic Lateral Sclerosis (ALS) and Frontotemporal dementia (FTD). This disease-causing expansion leads to a reduction in C9ORF72 expression levels in patients, suggesting haploinsufficiency could contribute to disease. To further understand the consequences of C9ORF72 deficiency *in vivo*, we generated a *c9orf72* mutant zebrafish line. Analysis of the spinal cord revealed no appreciable neurodegenerative pathology such as loss of motor neurons, or increased levels of neuroinflammation. However, detailed examination of *c9orf72^-/-^* retinas showed prominent neurodegenerative features, including a decrease in retinal thickness, gliosis, and an overall reduction in neurons of all subtypes. Structurally, analysis of rod and cone cells within the photoreceptor layer showed a disturbance in the outer cells of the retina and rhodopsin mis-localisation from rod outer segments to their cell bodies and synaptic endings. Thus, C9ORF72 may play a previously unappreciated role in retinal homeostasis and suggests C9ORF72 deficiency can induce tissue specific neuronal loss.

## Introduction

Hexanucleotide GGGGCC (G_4_C_2_) repeat expansions within the first intron of *C9ORF72* are now recognised as the most common genetic cause of the neurodegenerative disease spectrum Amyotrophic Lateral Sclerosis (ALS) and Frontotemporal dementia (FTD)^1,2^, accounting for ∼40% of familial ALS cases (∼10% sporadic) and ∼25% of familial FTD cases (∼6% sporadic) depending on the population analysed^3^. Furthermore, genetic variations at the *C9ORF72* locus are associated with sporadic ALS, as identified through large scale Genome Wide Association Studies which further link *C9ORF72* to sporadic ALS^4^. The number of G_4_C_2_ repeat expansions is highly variable between patients which can be as low as 30 repeats, rising up to 1000s of repeats in the most extreme cases^5,6^. However, *C9ORF72* repeat expansions are not always fully penetrant and repeat length does not necessarily correlate with disease severity^5,7,8^. How these repeat expansions cause neuronal loss in the ALS/FTD disease spectrum is still not completely understood, with several different mechanisms proposed which include both toxic gain of function (GoF) and loss of function (LoF).

GoF mechanisms includes both bi-directional transcription and translation of the pathogenic repeat expansion, leading to the accumulation of sense and antisense RNA foci, and dipeptide repeats (DPRs), respectively. RNA foci and DPRs have been detected in patient post mortem, and have been shown to be toxic in a variety of different *in vitro* and *in vivo* model systems^9–13^. Furthermore, the pathological G_4_C_2_ expansions lead to hypermethylation of C9ORF72 promoter thus leading to a reduction of *C9ORF72* transcript levels suggesting the potential implication of a LoF mechanism which could contribute towards neuronal loss. Indeed, a reduction in *C9ORF72* expression has been replicated over many different patient tissues and can be as pronounced as a 50% reduction in mRNA levels^10,14^. Despite this variety of different proposed mechanisms, the contribution of each to disease is unknown. Thus, the implications of both GoF and LoF must be studied in isolation to fully understand their consequences *in vivo*.

Although GoF mechanisms have been widely studied *in vivo*, LoF has been comparatively underexplored, with the function of *C9ORF72* still not completely understood^10^. *In vitro* studies suggest it to be part of the endosomal signalling network and to play a role in autophagy^15,16^. *In vivo*, knock-out (KO) mouse models suggest a role for *C9ORF72* in the immune system^10^ with KO mice robustly exhibiting enlarged spleens in conjunction with activation of macrophages and microglia^17,18^. Crucially *C9ORF72* deficiency does not cause neurodegeneration in mouse spinal cords^10,17,18^. Furthermore how C9ORF72 deficiency impacts other parts of the CNS, such as the retina, has yet to be explored *in vivo*, despite ocular defects being a feature of ALS patients^19–21^.

Despite many different KO mouse models being characterised in great detail (with respect to both their central nervous and lymphatic systems), *C9ORF72* deficiency has been poorly explored in other *in vivo* animal models^10^. Of note, within the neurodegeneration field, there are examples such as Parkinsons disease, where genetic mouse models have been shown to be resistant to neuronal loss compared to other vertebrate species such as rats and zebrafish^22–24^. Therefore, it is unknown whether *C9ORF72*^-/-^ mutant mice are simply impervious to neuronal loss or whether the latter is a general feature of vertebrate *C9ORF72* deficiency *in vivo*. Complementary investigations using alternative C9ORF72 deficient vertebrate model species would build a more comprehensive understanding of C9ORF72 function *in vivo*, but also to ascertain whether C9ORF72 deficiency can cause neuronal loss. Therefore, to fully understand the function of *C9ORF72 in vivo* and its contribution to ALS/FTD in vertebrates, characterising *C9ORF72* deficiency in additional model species is necessary. The zebrafish has shown to be a promising complementary *in vivo* vertebrate model of neurodegeneration, demonstrating the key markers of pathology, including neuronal loss and gliosis, even during early larval stages, in models of various neurodegenerative conditions^22,25–32^. Previous studies characterising *c9orf72* deficient larval zebrafish utilised a knock down approach with Morpholinos^33^, and indicated that knockdown of *c9orf72* in zebrafish resulted in axon outgrowth deficits in motor neurons and reduction in spontaneous movement^33^. However, adult *c9orf72* stable mutant zebrafish lines have yet to be described in detail, especially including at late-stage adult ages with respect to age, which is the largest risk factor for ALS/FTD in humans^16,34^.

Here we used CRISPR/Cas9 to generate a *c9orf72*-deficient zebrafish and characterise the spinal cord and retina at 24 months of age, when age-related neurodegeneration begins^35,36^. Similar to KO mice, *c9orf72*^-/-^ aged zebrafish do not show motor neuron loss within the spinal cord. Microglial activation was also not present within this region of the central nervous system (CNS). However, in depth analysis of the retina unexpectedly revealed prominent disruption of overall structure with a large decrease of retinal thickness within the aged mutants, which has been reported in ALS patients^37,38^. Most notably, GFAP immunoreactivity within the Müller glia (MG) was re-located to the apical end of the cell, demonstrating a gliosis-like phenotype. Microglia were also abnormally distributed within mutant retinas, congregating to the photoreceptor layer. Analysis of neurons within this layer revealed extensive photoreceptor outer segment disruption coupled with the ectopic localisation of rhodopsin in cell bodies and basal synaptic terminals. Furthermore, there were significant reductions in key neuronal subtypes throughout the retina. Analysis of all retinal markers at larval stages (5dpf) revealed no differences between WT and *c9orf72^-/-^*, suggesting the phenotypes observed here are due to progressive retinal degeneration in adults and not a developmental defect. Thus, the zebrafish represents a novel model to explore spontaneous neurodegeneration in a CNS tissue in C9ORF72 deficiency *in vivo*.

## Materials and Methods

### Zebrafish husbandry

All adult zebrafish were raised in standard conditions under project license PP5258250. The *c9orf72* mutant line was generated using CRISPR/Cas9 as previously described^39,40^ using standard Tracr RNA and a CrRNA targeting an *mwo1* restriction enzyme site within exon 2 of the zebrafish *c9orf72* (5’UAGAAGCUAAGCUCUGACUG), both purchased from Merck KGaA (Germany, Darmstadt). This complex was co-injected with Cas9 protein (M0386M, NEB, Ipswich USA) into the yolk of 1-cell stage WT zebrafish embryos. Once mutagenesis had been confirmed by restriction digest, F0 individuals were raised to adulthood and founders identified through crossing to WT. By using simple band shift on a standard agarose gel after PCR of the locus, a founder transmitting a large indel mutation to its F1 was identified. The *c9orf72* target locus was sequenced to confirm the mutation produced a frameshift, and then propagated to establish a working colony of *c9orf72* heterozygous individuals.

### Genotyping and QPCR validation

The *c9orf72* mutant line was genotyped used primers F 5’ CTTCGTCTTGGCTGAAAAGG and R 5’ TTGTAACCCTAGAAGAAAAACACAAA and genotypes assigned from simple band shift, due to the size of the mutation, on a 2% agarose gel. Transcript levels were determined by QPCR using primers for *c9orf72* (F 5’ GCTTCTACCTGCCTCTGCAC and R 5’ATCAACCTCCTCTGGCACAC) and normalised to the housekeeping gene *ef1a* (F 5’ TGGTACTTCTCAGGCTGACT and R 5’ TGACTCCAACGATCAGCTGT)^41^.

### Sample preparation and immunohistochemistry

All Zebrafish (WT or *c9orf72^-/-^*) were anesthetised by immersing into 0.02% aminobenzoic acid ethyl-methyl-ester [Sigma: MS222] in PBS followed by intracardial perfusion with PBS to remove residual blood and 4% PFA for fixation. Bodies were incubated in 4% PFA overnight for post-fixation. Spinal cord (SC) and skeletal muscle tissue were extracted using a stereo microscope. Skeletal muscle tissue was extracted from the right side of the dorsal part of the trunk, surrounding the spinal cord, starting caudal to the level of the operculum, until the level of the anterior border of the dorsal fin.

For 4c4 and GFAP immunofluorescent (IF) staining, SC was cryo-protected by incubation in 30% Sucrose, o/n, 4°C, followed by OCT embedding and cryo-sectioning (15mm). Sections were permeabilised with 0.25% Triton X-100, 10min, followed by blocking with 5% animal serum, 0.25% Triton X-100 in PBS for 1h, RT. Subsequently, primary antibodies were applied in blocking solution overnight at 4°C. The next day, sections were washed and incubated with appropriate fluorescent secondary antibodies for 1h, RT. Nuclei were counter stained with DAPI, followed by washing and mounting.

For ChAT and NMJ staining, SC and skeletal muscle tissue was embedded in 4% agarose and cut into 50mm or 150mm thick sections, respectively. IF staining of vibratome sections was performed according to a protocol adapted from Kusch *et al* 2012^42^. Vibratome sections were washed in (1) PBS-T (0.1% Triton X-100) for 10min, RT, (2) 50mM Glycine in PBS-T for 10min, RT, to remove residual PFA, and (3) in PBS-T for 15min, RT. Subsequently, sections were incubated in blocking solution (1% DMSO, 1% Donkey or Goat Serum, 1% BSA, 0.7% Triton X-100 in PBS) for 30min, RT. Primary antibodies were applied overnight at 4°C in blocking solution. The following day, sections were washed and incubated with secondary antibodies in blocking solution overnight, 4°C. The third day, nuclei were counterstained with DAPI for 10min, RT, followed by washing. Z-stacks of SC and muscle IF staining’s were acquired with the Zeiss Confocal 710.

For IHC of ocular tissue, Wild-type and mutant adult eyes were cryo-embedded in OCT prior to cryosectioning (18µm thickness). Sections were rehydrated by incubating the slides in PBS (1X) for 5 minutes at RT. Antigen retrieval was performed by heating the slides in 10mM sodium citrate, pH 6, for 20 mins. Once at room temperature, slides were washed three times in 0.1% PBS-Triton-X for 20 mins, prior to incubation in blocking solution (1% BSA, 10% goat serum in 0.1% PBS-Triton-X) at room temperature for 1 hour. Slides were then incubated with primary antibody solutions at 4°C overnight. The next day, excess antibodies were removed by washing slides three times in PBS for 20 minutes, followed by a 2-hour incubation with secondary antibodies at room temperature. Alexa-488 goat anti-rabbit and Alexa-647 goat anti-mouse secondary antibodies used were from Invitrogen (1:1000 dilution). Slides were washed three times in PBS for 20 minutes, the third wash containing DAPI stain (1:1000). Finally, slides were mounted with coverslips and dried in the dark overnight. Z-stack images in the central retina were acquired on the Zeiss 900 AiryScan 2 confocal microscope, using the 40x objective lens.

Information about the antibodies and dilutions used is summarised in Table 1.

**Table 1.** Antibodies.

| Antibody | Cat code | Company | Dilution | Species |
| --- | --- | --- | --- | --- |
| Glial fibrillary acidic protein (GFAP) | Z0334 | Dako | 1:500 | Rabbit |
| 4c4 | - | Generated in house | 1:100 | Mouse |
| Choline acetyl transferase (ChAT) | AB144P | Sigma-Aldrich | 1:100 | Goat |
| Synaptic vesicle glycoprotein 2A (SV2A) | SV2 | DSHB | 1:100 | Mouse (IgG1) |
| Donkey anti mouse IgG (H+L) Cy3 | 715-165-150 | Jackson | 1:200 | Donkey |
| Donkey anti rabbit 647 | A31573 | ThermoFisher | 1:1000 | Donkey |
| Donkey anti goat 555 | A21432 | ThermoFisher | 1:500 | Donkey |
| Goat anti mouse IgG1 488 | A21121 | ThermoFisher | 1:1000 | Goat |
| Bungarotoxin | B35451 | ThermoFisher | 1:1000 | - |
| ZRF-1 (GFAP) | AB_10013806 | ZIRC | 1:200 | Mouse |
| Ribeye A |  | Gift from T. Nicholson | 1:2000 | Rabbit |
| GNAT2 | PM075 | MBL | 1:100 | Rabbit |
| PKC-α | Sc-208 | Santa Cruz | 1:300 | Rabbit |
| HuC/D | A21271 | Invitrogen | 1:300 | Mouse |
| GS | 66323-1-Ig | Proteintech | 1:200 | Mouse |
| ZPR-3 | AB_10013805 | ZIRC | 1:200 | Mouse |
| ZPR-1 | AB_10013803 | ZIRC | 1:200 | Mouse |
| Rhodopsin (Rho) | n/a | Gift from David Hyde | 1:2000 | Rabbit |
| Zo-1 | 33-9100 | Invitrogen | 1:100 | Mouse |
| Goat anti rabbit-Alexa Fluor 488 | A11008 | Invitrogen | 1:1000 | Goat |
| Goat anti mouse-Alexa Fluor 647 | A21235 | Invitrogen | 1:1000 | Goat |

### Image analysis

Maximum projections of Z-stacks were thresholded, converted into binary mask and measured in Fiji/ImageJ. Threshold (4c4, GFAP, BF) was chosen manually for each batch which was processed simultaneously for WT and *c9orf72^-/-^* fish. Microglia and astrocyte area measurements were calculated as percentage of area post-thresholding normalised to BF area of the whole spinal cord section. Microglia, ChAT and NMJ counts were counted manually per section. All image processing was performed blinded. Statistical analysis was performed as unpaired, parametric two-tailed t-test.

All retinal quantifications were performed using Fiji/Image J on 3D z-stacks. Neuronal numbers were determined by manually counting cells positive for both the neuronal markers of interest and DAPI, within a 100×100×10μm ROI using the ‘multi-point’ tool. 4c4^+^ microglia were counted in the entire field of view, and in individual retinal layers. Retinal thickness measurements were carried out in DAPI stained sections, and individual layer measurements were normalised to the overall retinal thickness from the top of the INL to the bottom of the RGC layer and expressed as a proportion of overall retinal thickness. Following Shapiro-Wilk normality tests to confirm normal distribution, statistical significance was determined using unpaired two-tailed t-tests when comparing a single measurement between two groups. Two-way ANOVA was used when comparing multiple measurements within one image, e.g. different layers in the same retina. GFAP immunofluorescence distribution along the apicobasal axis was quantified using IMARIS. Briefly, Gfap staining was segmented by generating a surface on IMARIS and split into the apical and basal halves. The volume was quantified automatically on IMARIS for the apical and basal regions, and a ratio between the two was quantified to calculate the apicobasal distribution in WTs and mutants.

## Results

### Mutant generation and validation of the *c9orf72^-/-^* mutant zebrafish line

A single *C9ORF72* orthologue was identified in zebrafish (ENSDARG00000011837) using ENSEMBL. The zebrafish orthologue possesses ∼75% protein homology to the human protein as well as a similar exonic structure and conserved synteny (Supplemental Figure 1). To further understand the consequences of *c9orf72* deficiency *in vivo* we generated a loss of function zebrafish line by targeting exon 2 using CRISPR/Cas9 (Figure 1A). The selected *c9orf72* mutant allele contained a combined 5bp deletion and 90bp insertion, forming a frameshift and the generation of a premature stop coding within exon 2 (Figure 1A-C). Due to the lack of reliable antibodies for *c9orf72* in zebrafish we measured *c9orf72* transcript levels in the homozygous mutants at 5dpf. This revealed a 66% reduction of *c9orf72* mRNA (Figure 1D, p=0.011, n=4, Welch’s two tailed t-test) in the *c9orf72^-/-^* compared to WT controls demonstrating the frameshift allele causes nonsense mediated decay, indicating the mutation is loss of function.

**Figure 1.**
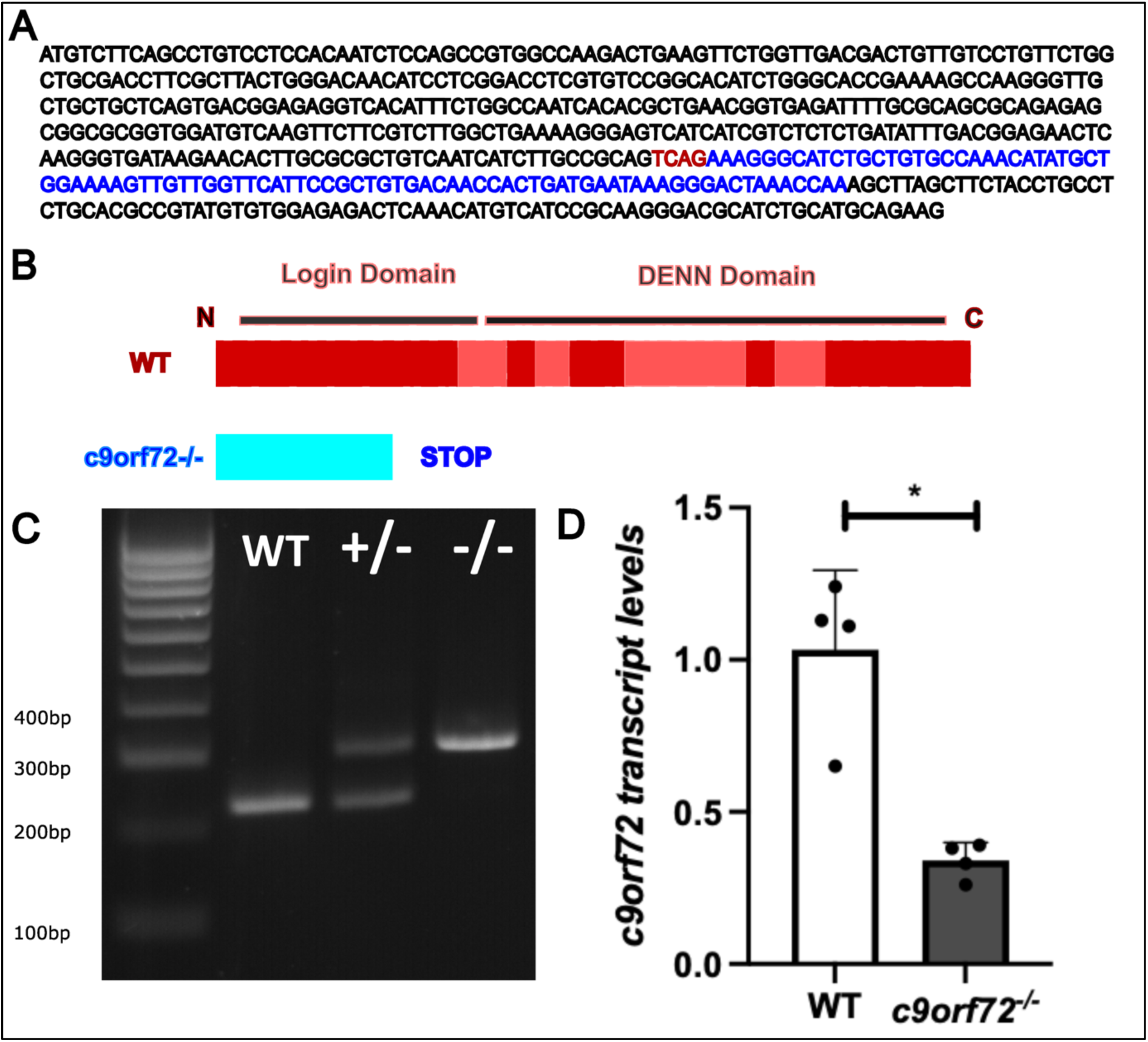
Characterisation of the *c9orf72* loss of function mutation. A) Exon 2 of *c9orf72* illustrating the frameshift mutation generated through CRISPR/Cas9. The frame shift allele is composed of a 5bp deletion (red script) and 90bp insertion (blue script) within WT exon 2 (black script). B) The frameshift mutation when translated results in a premature stop codon within the login domain, deleting the majority of the protein when compared to the WT. Different shades of red represent alternating exons within the WT protein. C) Example genotyping gel demonstrating the large in-del mutation visible by gel electrophoresis. D) The mutation leads to a large 66% reduction in *c9orf72* transcript levels in the *c9orf72^-/-^* (p=0.011, n=4, Welch’s two tailed t test), demonstrating activation of nonsense mediated decay by the frameshift allele, indicative of loss of function.

### Absence of neuronal loss and NMJ architecture in the spinal cord of *c9orf72^-/-^*

ALS is characterised by the loss of motor neurons (MNs), leading to muscle weakness and ultimately death due to respiratory failure^43^. Crucially, the neuromuscular junction (NMJ), the synaptic connection between MNs and skeletal muscle, is subject to degeneration early during ALS pathology which precedes MN loss. As such, we were interested in understanding the effect of *C9ORF72* deficiency on spinal cord MNs and NMJs in adults at an advanced stage of their life course. Zebrafish are known to have an average lifespan up to 42 months^44^, and age-related neurodegeneration begins to occur at approximately 24 months^35,36^, as such we selected 24 month of age for analysis of *c9orf72^-/-^* mutants. Firstly, we stained for the mature cholinergic MN marker choline acetyl transferase (ChAT). No difference in ChAT^+^ MN numbers was observed between WT (2.92 ChAT^+^ MNs per section) and *c9orf72*^-/-^ (3.37 ChAT^+^ MNs per section; p=0.038, n=15), highlighting no MN loss (Figure. 2A-C). NMJ degeneration precedes MN loss in ALS mouse models as well as clinical studies ^45–47^ therefore we analysed NMJ structure within our *c9orf72^-/-^*to potentially detect subtle changes towards motor neuron health. Skeletal muscle was stained for synaptic vesicle 2 (SV2, pre-synaptic marker) and acetylcholine receptor (AChR, post-synaptic marker, labelled with bungarotoxin (Btx), Figure. 2D-E). We distinguished between SV2^+^/Btx^+^ (innervated NMJ), SV2^+^/Btx^-^ (pre-synapse only) and SV2^-^/Btx^+^ (post-synapse only) (Figure. 2D-E). However, no difference in any of the NMJ subtypes was observed (Figure. 2D-E). Taken together, *c9orf72* deficiency in adult zebrafish does not lead to spinal cord MN loss or NMJ degeneration at 24 months of age.

**Figure 2.**
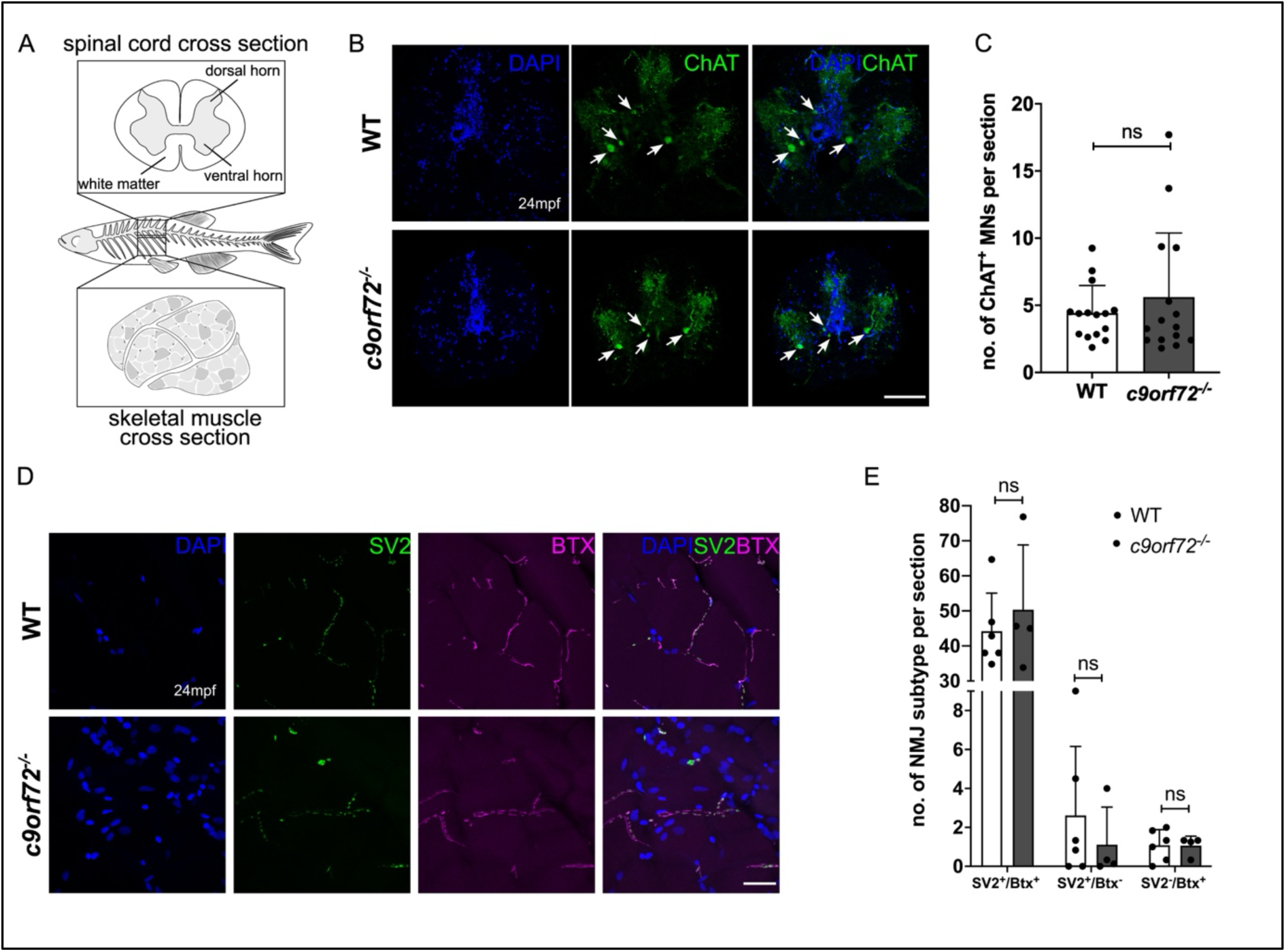
*c9orf72* deficiency does not lead to MN loss or NMJ degeneration in adult zebrafish spinal cord. A) Schematic diagram of adult zebrafish, highlighting the spinal cord region used for analysis. B) Representative immunofluorescent image of ChAT staining in WT and *c9orf72*^-/-^ spinal cord of 24-month-old zebrafish. Scale bar: 100μm. C) Quantification of A. MN numbers were not significantly different between WT (2.92 ChAT^+^ MNs per section) and *c9orf72^-/-^*(3.37 ChAT^+^ MNs per section) zebrafish. Unpaired, two-tailed and parametric t-test, p=0.038, n=15 per genotype. D) Representative immunofluorescent image of NMJ staining in WT and *c9orf72^-/-^* muscle of 24-month-old zebrafish. SV2: pre-synaptic marker; Btx: post-synaptic marker labelling AChRs. Scale bar: 25μm. E) Quantification of D. NMJ integrity was not altered in *c9orf72*^-/-^ zebrafish. SV2^+^/Btx^+^: WT=44.19, *c9orf72^-/-^*=50.32, p=0.52; SV2^+^/Btx^-^: WT=2.61, *c9orf72^-/-^*=1.12, p=0.47; SV2^-^/Btx^+^: WT=1.08, *c9orf72^-/-^* =1.06, p=0.96. Unpaired, two-tailed and parametric t-test, n-WT:6, n-*c9orf72^-/-^*:4.

### Lack of gliosis in the spinal cord *c9orf72^-/-^* mutants

Consistent with our findings, *C9ORF72* KO mouse models also do not exhibit MN loss or NMJ degeneration, but do develop an increased neuroinflammatory signature, including microgliosis ^18,48^. Therefore, we were interested in understanding if glial activation was present in the CNS of our *c9orf72^-/-^* zebrafish. Spinal cord sections were stained for the zebrafish specific microglial marker 4c4^49^ and glial fibrillary acidic protein (GFAP), labelling astrocytes and radial glia. Microglial number was not significantly altered in the spinal cord of *c9orf72^-/-^* zebrafish (41.85 microglia per section) compared to WT (43.32 microglia per section, p=0.83, n=13-14, Figure. 3A and B). Furthermore, microglial area was also not notably increased in *c9orf72^-/-^* (WT: 2.9%, *c9orf72^-/-^*: 3.37%, p=0.50, n=13-14, data not shown). Equally, GFAP^+^ area within the spinal cord was not significantly altered comparing WT (20.34%) and *c9orf72*^-/-^ (25.18%, p=0.18, n=13-14, Figure. 3C and D). Overall, these results suggest the absence of neuroinflammation in adult *c9orf72^-/-^*zebrafish spinal cord. To identify potentially more subtle neuroinflammatory signatures within the CNS, we analysed chitotriosidase activity levels (a marker of generalised inflammation and potential ALS biomarker^50^) in brain homogenates. No statistical differences were found between mutant and WT confirming a lack of inflammatory signatures in the *c9orf72^-/-^*(Supplemental Figure 2A). To ascertain whether *c9orf72* deficiency results in lysosomal dysfunction within the CNS, we analysed the activity of the lysosomal enzymes, β-hexosaminidase and β-galactosidase in brain homogenates. The activity of both enzymes was not significantly altered between genotypes (Supplemental Figure 2B and C, n=3-4), suggesting lysosomal function is not grossly altered in the brains of our *c9orf72^-/-^*.

**Figure 3.**
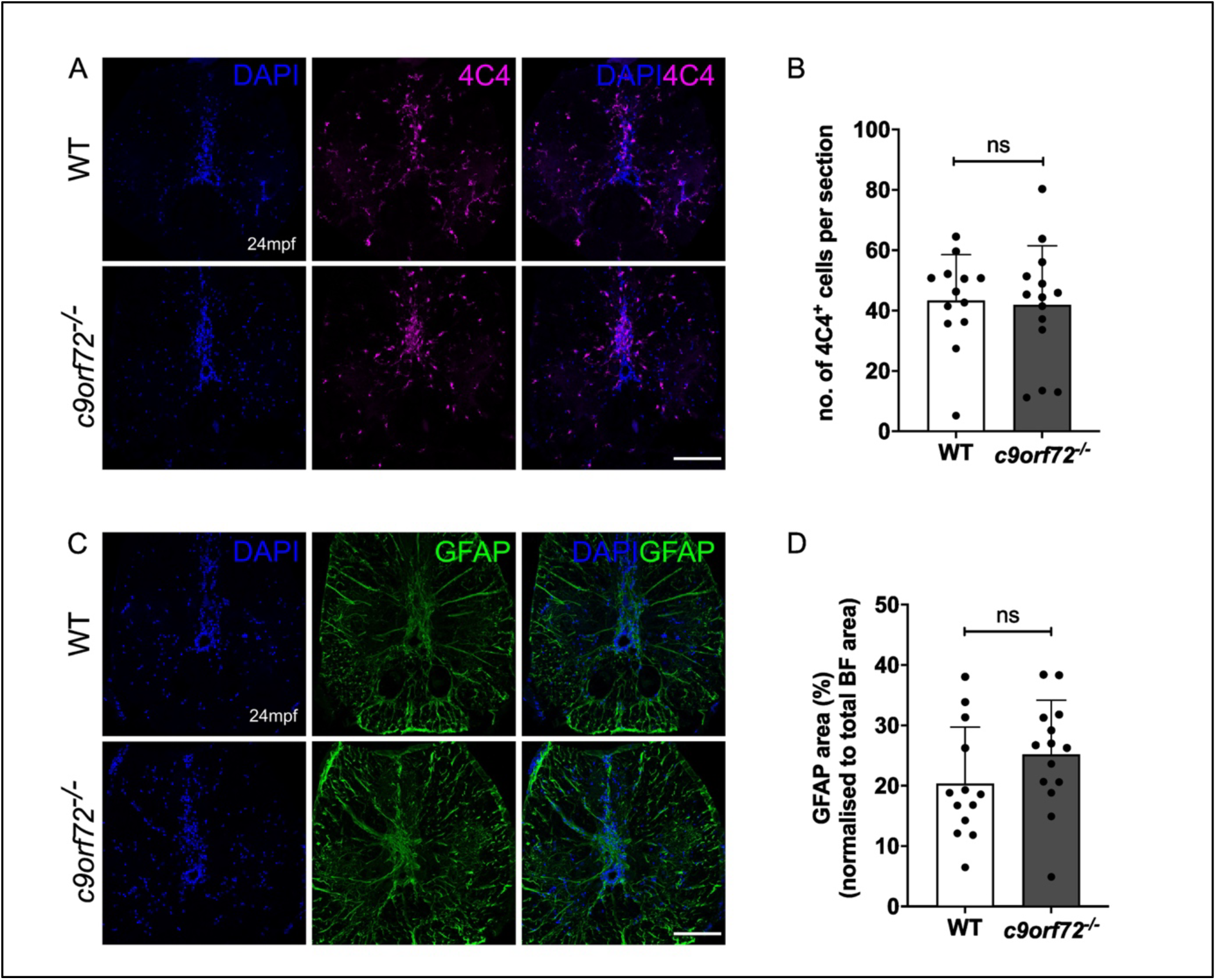
*c9orf72*^-/-^ adult zebrafish do not exhibit micro– or astrogliosis. A) Representative immunofluorescent image of 4c4^+^ microglia in WT and *c9orf72*^-/-^ spinal cord of 24-month-old zebrafish. Scale bar: 100μm. B) Quantification of A. Microglial numbers per spinal cord section are unaltered between WT (43.32 4c4^+^ microglia) and *c9orf72^-/-^* (41.85 4c4^+^ microglia). Unpaired, two-tailed and parametric t-test, p=0.83, n-WT:13, n– *c9orf72^-/-^*:14. C) Representative immunofluorescent image of GFAP staining in WT and *c9orf72*^-/-^ spinal cord of 24-month-old zebrafish. Scale bar: 100μm. D) Quantification of C. GFAP^+^ area within spinal cord, normalised to spinal cord size, does not differ between WT (20.34%) and *c9orf72^-/-^* (25.18%). Unpaired, two-tailed and parametric t-test, p=0.18, n-WT:13, n-*c9orf72^-/-^*:14.

### *c9orf72*^-/-^ mutants show signs of neurodegeneration in the retina

Ocular defects in *C9ORF72*-associated disease have been described in patients^37,38^, as well as in *Drosophila* models of c9orf72-associated ALS^51^. However, how C9ORF72 deficiency impacts the retina has yet to be described in a vertebrate model system. The retina is a highly structured and conserved CNS tissue, organised into distinct nuclear layers, separated by synaptic plexiform layers. Here we focused on the central retina (Figure 4A), as it has been shown to display retinal degeneration earlier than peripheral regions in the ageing WT zebrafish^52,53^.

**Figure 4.**
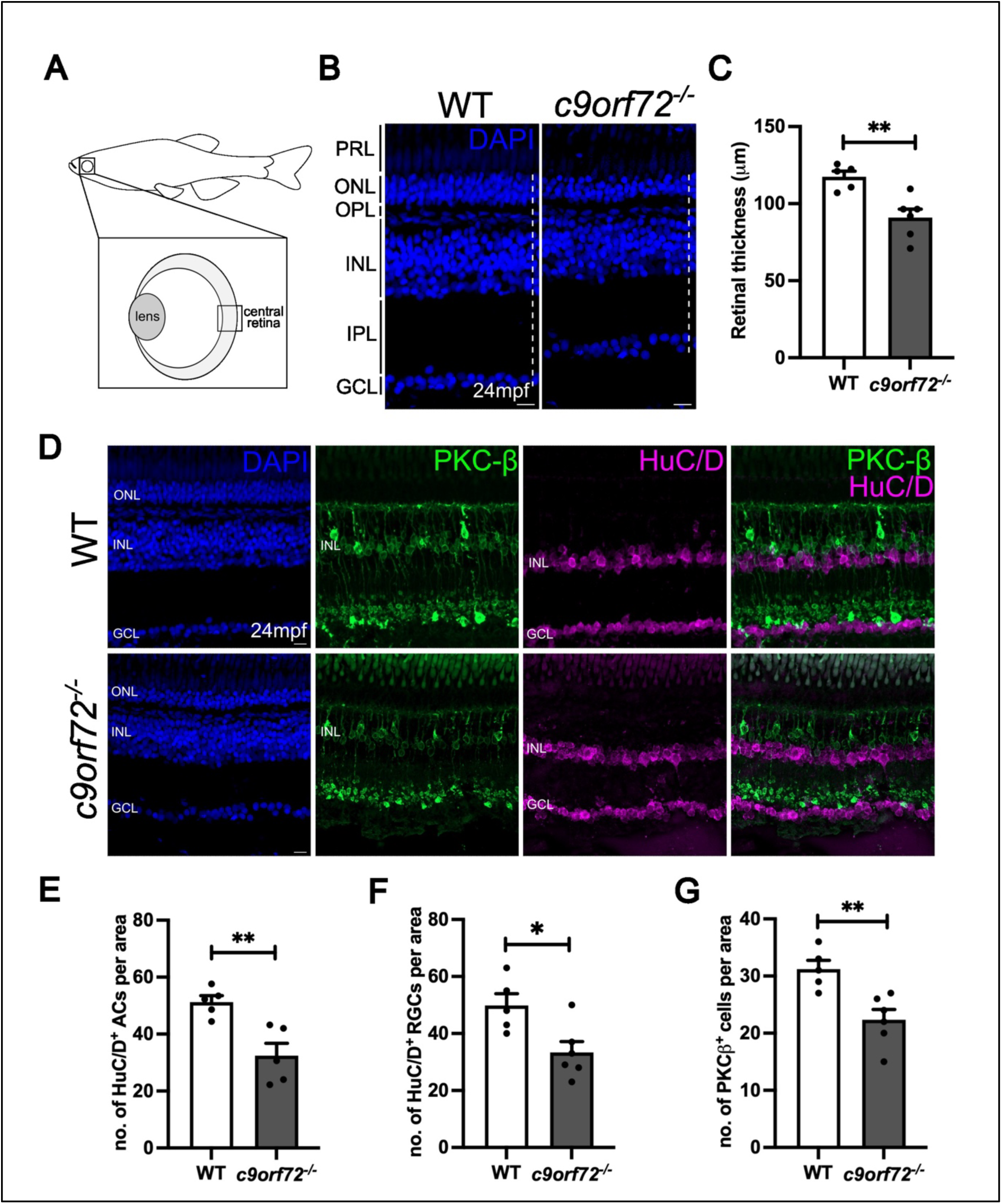
*c9orf72^-/-^* mutants exhibit signs of degeneration of inner retinal neurons. A) Schematic diagram of adult zebrafish retina, highlighting the central retinal region used for analysis. B) DAPI (blue) staining showing the nuclear layers of the WT and *c9orf72^-/-^* retina. In the innermost retina, retinal ganglion cell (RGC) somata make up the ganglion cell layer (GCL) and project processes into the inner plexiform layer, where they form synapses with bipolar and amacrine cells, the cell bodies of which are found in the inner nuclear layer (INL). Apical to the INL is the outer plexiform layer (OPL), where bipolar cells and horizontal cells connect to the photoreceptors in the outer nuclear layer (ONL). In the outermost retina, the inner and outer segments of rod and cone photoreceptors make up the photoreceptor layer (PRL). Dotted line represents the approach to measure retinal thickness. C)Quantification of mean retinal thickness of WT and *c9orf72^-/-^*, unpaired t-test; p=0.0041. D) Antibody staining for bipolar cell marker, (PKC-β, green), amacrine and retinal ganglion cells (HuC/D, magenta) and nuclei stained with DAPI (blue) in WT and *c9orf72^-/-^* retinas. E) Quantification of number of HuC/D^+^ amacrine cells in INL per 100μm x 100μm ROI; unpaired t-test; p=0.0051. F) Quantification of number of HuC/D^+^ retinal ganglion cells in GCL per 100μm x 100μm ROI; unpaired t-test; p= 0.0176. G) Quantification of number of PKC-β^+^ bipolar cells in INL per 100μm x 100μm ROI; unpaired t-test; p= 0.0055 .n=5-6 retinas per genotype; Scale bars, 50μm.

As thinning of the retina is a hallmark sign of neurodegeneration, we assessed overall tissue thickness in sections of WT and *c9orf72*^-/-^ mutant retinas. The mean thickness of *c9orf72^-/-^* retina was approximately 20% thinner than its WT control at 24 months (WT: 117.4 ± 3.6 μm *c9orf72^-/-^*: 90.9 ± 5.5 μm) (Figure 4B and C). Measurements of the thickness of individual layers did not reveal any significant differences between WT and *c9orf72^-/-^* retinas (data not shown), suggesting global degeneration at each neuronal layer. Next, to determine whether the observed thinning was due to the loss of a specific neuronal cell type, we performed immunostaining for known neuronal markers in the inner retina of WT and *c9orf72^-/-^* (Figure 4D). Antibody staining for HuC/D, a marker for amacrine and retinal ganglion cells, revealed a 36.8% (p=0.0051) and 33.0% (p=0.0176) reduction in numbers, respectively (Figure 4E and F). Similarly, we observed 28.4% fewer (p=0.0055) PKC-β positive bipolar cells, another type of retinal interneuron, in *c9orf72^-/-^* mutants compared to WT controls (Figure 4G). Retinal thickness and neuronal markers were also investigated at 5 days post fertilisation (5dpf), However no differences were found between WT and *c9orf72^-/-^* mutants at this earlier timepoint, indicating that retinal development is unperturbed (Supplemental Figure 3). Together, these data identify retinal degeneration phenotypes in 24 month old *c9orf72^-/-^* mutants and suggest that the retinal thinning observed may be attributed to an overall loss of neurons in the INL (inner nuclear layer, approx. 30% of each cell type), rather than a cell-type specific degeneration.

### *c9orf72*^-/-^ mutants exhibit signs of photoreceptor degeneration

Having identified signs of neurodegeneration in inner retinal neurons of *c9orf72^-/-^* mutants, we next examined the consequences of *c9orf72* deficiency on the outer retinal cell-types. Quantification of mean cell numbers showed a ∼23% decrease in the number of all cone photoreceptors in *c9orf72* mutants compared to WTs, as labelled by the pan-cone marker Gnat2 (Figure 5A and B). Similarly, staining with Zpr-1 revealed a decrease in cell numbers (Figure 5A and C). In addition, the number of Zpr-3-positive rod and green cone outer segments was also decreased in the *c9orf72* mutants (Figure 5D-F), as well as the rod photoreceptors as observed by rhodopsin (Rho) antibody labelling (Figure 5D and G). together confirming that the loss of photoreceptors was not sub-type specific. Thus, similarly to the inner retina, we observed a global loss of all neuronal populations of photoreceptors in the *c9orf72^-/-^* retina. Strikingly, aside from the presence of fewer rods in the aged *c9orf72^-/-^* retina (Figure 5F), we observed the mis-localisation of rhodopsin to the cell bodies and processes in the OPL (outer plexiform layer), instead of being confined to the outer segments as observed in WTs (Figure 5E). Although the occurrence of this was sporadic, with only 1-3 cells per central retinal region analysed, this phenomenon was exclusive to the *c9orf72^-/-^*, and not seen in any WT controls. Furthermore, investigation of Gnat2, Zpr-1, Rho and Zpr-3 demonstrated no differences between genotypes at 5dpf suggesting again the dysfunction exhibited at the 24 month time point in *c9orf72^-/-^* is not developmental in origin (Supplemental Figure 4).

**Figure 5.**
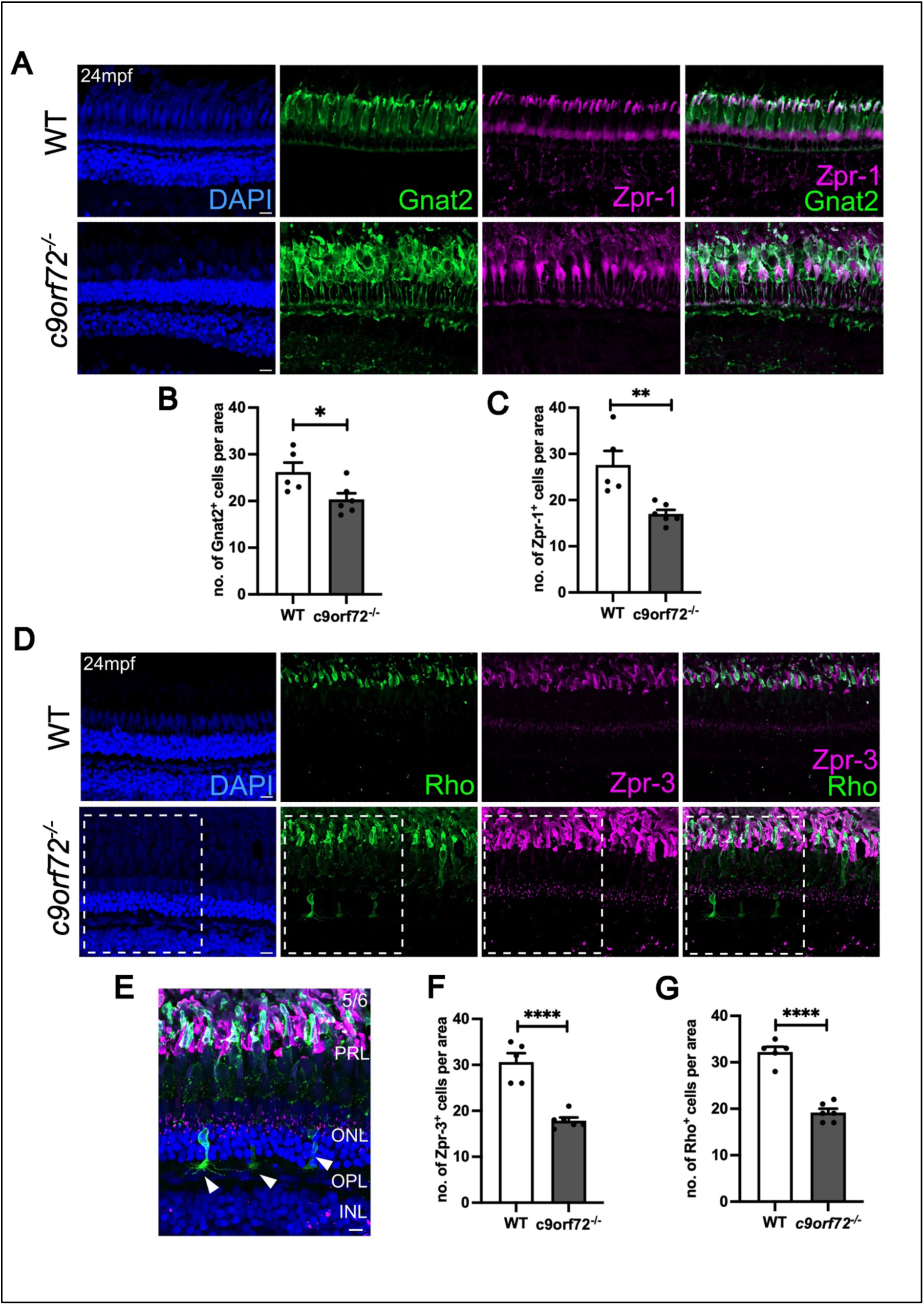
Photoreceptor degeneration in *c9orf72^-/-^*mutants. A) Immunostaining for pan-cone marker, Gnat2 (green) and double cone marker, Zpr-1 (magenta) in WT and c9orf72-deficient retinal cryosections. Nuclei labelled with DAPI (blue). B) Quantification of mean number of Gnat2-positive cone photoreceptors in WT and *c9orf72*-deficient retinas;100μm x 100µm x 10 μm ROI; unpaired t-test, p= 0.0333. C) Quantification of mean number of Zpr-1-positive cone photoreceptors in WT and *c9orf72*-deficient retinas;100μm x 100μm x 10µm ROI; unpaired t-test, p= 0.0055. D) Antibody staining for rod photoreceptor marker rhodopsin (Rho, green) and rod and cone photoreceptor marker (Zpr-3, magenta) and nuclei stained with DAPI (blue) in WT and *c9orf72^-/-^* retinas. E) Close up of *c9orf72^-/-^* retina from (D) with nuclei labelled with DAPI (blue). F) Quantification of mean number of Zpr-3^+^ rod and short double cone photoreceptors in WT and c9orf72-deficient retinas;100μm x 100μm ROI; unpaired t-test, p<0.0001. G) Quantification of number of Rho^+^ rods in ONL per 100μm x 100μm x 10µm ROI; unpaired t-test; p <0.0001; white arrowheads indicate displaced photoreceptors; phenotype observed in 0/5 WT retinas and 5/6 *c9orf72^-/-^* retinas; n=5-6 retinas per genotype; PRL, photoreceptor layer; ONL, outer nuclear layer; OPL, outer plexiform layer; INL, inner nuclear layer. Scale bars, 10μm.

### Gliosis and microglial redistribution accompany neuronal degeneration in the c*9orf72^-/-^* mutant retinas

As we observed neurodegeneration in the retina we asked whether there were also signs of glial activation ^54,55^, similar to what is observed in the ageing retina^52^. To investigate whether *c9orf72* deficiency impacts retinal glia we performed immunostaining for markers MG, the principal macroglia of the retina, as well as microglia. A characteristic sign of MG gliosis is the upregulation and apical redistribution of Gfap^55–57^. In WT retinas, Gfap immunoreactivity was confined predominantly to the basal end-feet of the MG (Figure 6A, asterisk). In contrast, in *c9orf72^-/-^* mutants, we observed Gfap staining extending towards the photoreceptors in the outer retina (Figure 6A, arrowhead). Quantification of apicobasal distribution of Gfap showed that relative to WT controls, the *c9orf72^-/-^* mutants showed an increase in Gfap staining in the apical half of the retina (Figure 6B and Supplemental Figure 5). Next, to determine whether C9orf72-deficient retinas exhibit signs of inflammation, we investigated the localisation, morphology and numbers of microglia using immunohistochemical approaches (Figure 6C and D). The average number of 4c4-positive microglia was not significantly different between WT and *c9orf72^-/-^* mutants (Figure 6D). However, the proportion of microglia located in the outer retina versus the basal regions was 9-fold higher (p=0.0035) in *c9orf72^-/-^* mutants compared to WT controls, consistent with the photoreceptor degeneration phenotypes (Figure 6E). However, we did not detect 4c4-positive cells in close proximity to the displaced ID4 photoreceptors (data not shown) or obvious morphological signs of microglial activation (i.e. ramified vs spheroid) in *c9orf72^-/-^* mutant retinas. Thus, we observed signs of glial activation in MG and microglia in the absence of C9ORF72 function, consistent with a general degeneration phenotype in the retina. In a similar fashion to the neuronal markers, there were no differences between genotypes in glial markers at 5dpf (Supplemental Figure 6).

**Figure 6.**
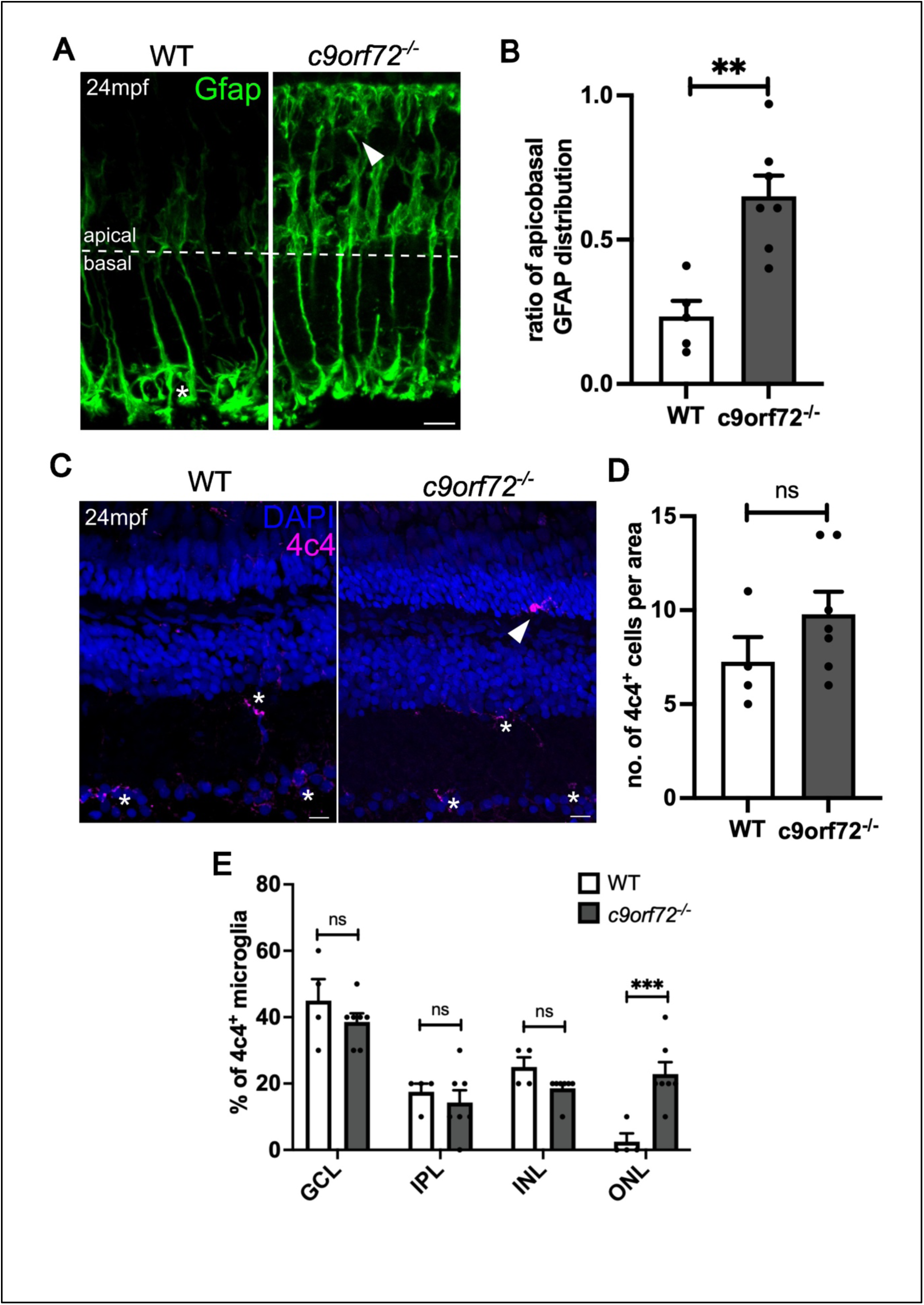
Gliosis phenotypes in *c9orf72^-/-^* deficient retinas. A) Immunostaining for gliosis marker, Gfap (green) in WT and *c9orf72*-deficient retinal cryosections. Nuclei labelled with DAPI (blue). B) Ratio of mean Gfap distribution in apical versus basal regions of the retina; unpaired t-test, p= 0.0016. C) Antibody labelling of retinal microglia with 4c4 (magenta), cell bodies labelled with DAPI (blue) D) Quantification of the average number of 4c4^+^ microglia per image; unpaired t-test; p= 0.2096. E) Quantification of the proportion of 4c4^+^ microglia observed in the ganglion cell layer (GCL), inner plexiform layer (IPL), inner nuclear layer (INL), and the outer nuclear layer (ONL); Two-way ANOVA, Šidâk’s multiple comparisons test; GCL, p=0.5932; IPL, p=0.9466; INL, p=0.5932; ONL, p=0.0008. n=5, WT retinas; n=6, *c9orf72^-/-^*retinas; Scale bars, 10μm.

## Discussion

*C9ORF72* hexanucleotide repeat expansions are the most common cause of both familiar and sporadic ALS/FTD. Nevertheless, it is unclear if *C9ORF72* LoF, DPR/RNA foci GoF, or a synergy of both is causative for disease. Here, we have established a stable *c9orf72* deficient zebrafish line to understand the impact of *C9ORF72* LoF in an adult vertebrate *in vivo* system. Our LoF *c9orf72^-/^*^-^ mutants had conventional development and lived for 24 months without any overt signs of ill health. Neurodegeneration was restricted to the retina and comparatively absent from the spinal cord, suggesting that *C9ORF72* deficiency alone is not sufficient to induce a neurodegenerative phenotype in the spinal cord, as demonstrated from mammalian studies^18,58^.

### Comparison with other C9ORF72 deficient models

Contrary to our findings, previous reports in the field described *c9orf72* deficient zebrafish models with comparatively severe phenotypes of axonopathy and early larval lethality^33,59^. However, both of these models were based on RNA interference which can have potential toxic off target effects^60,61^. These toxic side effects classically present as neuronal apoptosis, and thus neurodegenerative studies in zebrafish utilising RNA interference are difficult to interpret, unless validated with a traditional loss of function stable mutant^61,62^. Furthermore, the discrepancy between stable mutant and RNAi phenotypes is unlikely due to our mutant allele being hypomorphic. Our stable mutation is present within an early exon of *c9orf72*’s single transcript, ablates a key domain and activates nonsense mediated decay of the mutant mRNA. As zebrafish *c9orf72* lacks any direct paralogues, its inhibition should not activate the transcriptional adaptation pathways which has previously explained phenotypic discrepancies between the RNA interference based models and their respective stable loss of function mutants^63,64^. Additional traditional CRISPR/Cas9 loss of function *c9orf72* zebrafish mutants have also been reported without any accounts of impacted viability^65,66^. Unexpectedly, our aged mutants did not exhibit similar microglial activation and an inflammatory profile in the wider CNS as described in mouse models^10,17,18^. This may be due to a fundamental difference in microglial function between mammals and fish^67^.

### Retinal Pathology and C9ORF72

Given the lack of neuroinflammation within the wider CNS, the gliosis, neuronal loss and degeneration within the retinal tissue of our aged zebrafish *c9orf72^-/-^* was unexpected. Studies in other vertebrate models of C9ORF72 deficiency did not characterise the retina, as such we do not know if they also develop retinal phenotypes. It will be highly prudent to determine how common retinal degeneration is within other C9ORF72 deficient vertebrate models, such as the mouse or rat^10,58^ and also in patients harbouring *C9ORF72* repeat expansion mutations in order to determine whether retinal pathology is a universal feature of *C9ORF72* loss of function. The retinal phenotypes show an unexpected role for *C9ORF72* in homeostasis of the retina and demonstrate that C9ORF72 deficiency alone can induce spontaneous neurodegeneration, which is relevant for *C9ORF72* linked ALS/FTD. Of note, TDP43 pathology, as well as C9ORF72 specific DPR pathology have been described within the retina of *C9ORF72* linked ALS/FTD patients^38,68^. In the latter, each pathology is found in distinct areas of the retina (OPL and INL, respectively), in a similar fashion to the wider CNS where TDP43 and DPR aggregates are not found in the same neurons^68,69^. Several other ALS linked genes are also associated with the neurodegenerative eye diseases^70–74^. *TBK1, OPTN* and *ATXN2* variants all cause familial ALS but also glaucoma^70–74^. Furthermore, variants in *NEK1* and *C21ORF2* are causative for both ALS and retinitis pigmentosa^73,75,76^. However, C9ORF72’s precise function in retinal neurons is currently not clear.

We identified basally-translocated Rho staining in rod photoreceptors in the *c9orf72^-/-^* retina^77^. This mislocalisation from outer segments to the cell body and neurites of these cells points to a potential involvement of C9orf72 in opsin trafficking. Indeed, similar rhodopsin mis-trafficking phenotypes were reported in mice lacking unconventional Myosin 1C (MYO1C), a motor protein, as well as in Centrosomal Protein 290 (Cep290)-deficient zebrafish, leading to rhodopsin mislocalisation to the inner segment and cell bodies^78,79^. Given the reported interactions between C9ORF72 and Rab proteins, which can influence motor proteins involved in intracellular trafficking^80^, it is plausible that C9ORF72 is required for the process of intracellular trafficking, including that of photoreceptor opsins. Cep290, a ciliary protein present in the cilium connecting the inner and outer segments of photoreceptors is essential for the intracellular trafficking between these subcellular compartments, again pointing to a role of C9orf72 in these events. Future studies will be devoted to establishing how early retinal phenotypes are evident in our zebrafish *c9orf72* mutants and exploring the role of C9orf72 in motor protein function. This will be important for establishing whether early retinal changes can be used as an early indicator of neurodegeneration in the wider CNS.

Retinal thinning is a key feature of our *c9orf72* mutant zebrafish, which is also present in ALS patients that do not otherwise present with ophthalmic disease^19–21^. Ocular assessment is currently absent from the diagnostic pipeline in clinics, and hence confirming the presence of similar retinal phenotypes in mammalian model organisms may provide critical evidence for the need for thorough ocular tests as part of ALS diagnosis. As the retina is part of the CNS, most neurodegenerative diseases that affect neurons in the brain or spinal cord also impact retinal neurons. For example, retinal thinning and neuronal loss are common features presented by Alzheimer’s disease (AD) patients^81^ even in early stages of the disease and despite its progressive nature. The same is also true for Huntington’s disease^82^, further highlighting the potential usefulness of retinal assessment for the diagnosis of other neurodegenerative diseases^83–85^.

## Conclusions

Our study shows novel pathology in the context of *c9orf72* deficiency *in vivo*, namely retinal neurodegeneration, demonstrating *c9orf72* LoF can trigger neuronal loss spontaneously. Additionally, we have characterised a stable *c9orf72*^-/-^ zebrafish, which is highly valuable to understand not only the mechanism of retinal degeneration we have described, but also to study synergistic effects of ALS risk factors, mutations, and pathways potentially causative of neurodegenerative disease in a rapid and cost-effective *in vivo* vertebrate model system.

## Acknowledgements

We thank Prof Siddharthan Chandran and Dr Bhuvaneish T. Selvaraj for their mentorship and feedback during the generation of this manuscript.

## Funding

RBM – BBSRC David Phillips Fellowship (BB/S010386/1) and Moorfields Eye Charity PhD Studentship (GR001148).

Andrea Salzinger is supported by a Marie Skłodowska-Curie ETN grant under the European Union’s Horizon 2020 Research and Innovation Programme (Grant Agreement No 860035 SAND).

### Author Contributions

Conducted and analysed experiments: AS, NJ, TT, MK

Experimental design: AS, NJ, TMT, MK, DL, CB, TB, RM

AS, NJ, RM, MK wrote the manuscript.

All authors read and approved the final manuscript.

