## Supplemental Figures for "C9ORF72 deficiency results in degeneration of the zebrafish retina *in vivo*"

### Supplementary Figures

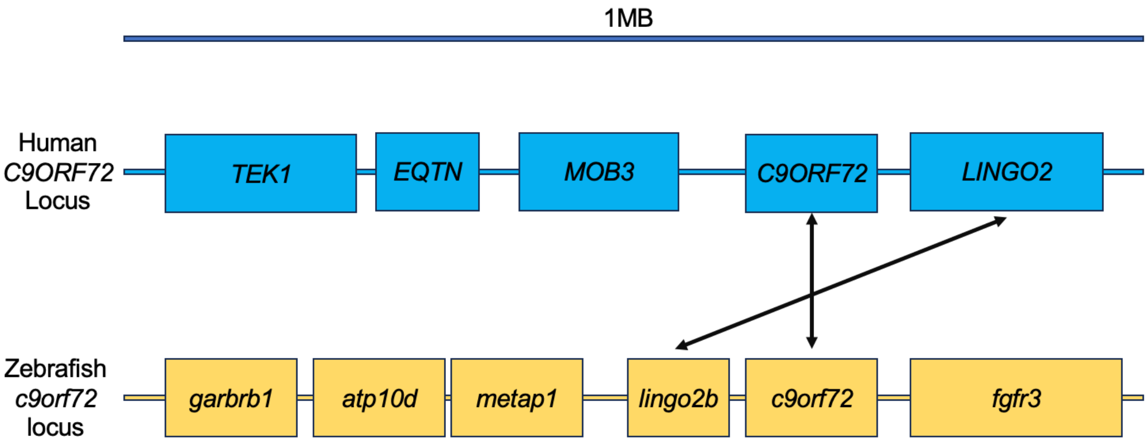


**Supplemental Figure 1.** Schematic of conserved gene synteny between the human *C9ORF72* and zebrafish *c9orf72*. In both species the genes encoding C9ORF72 and LINGO2 are located within 0.5mb of each other on the same chromosome.

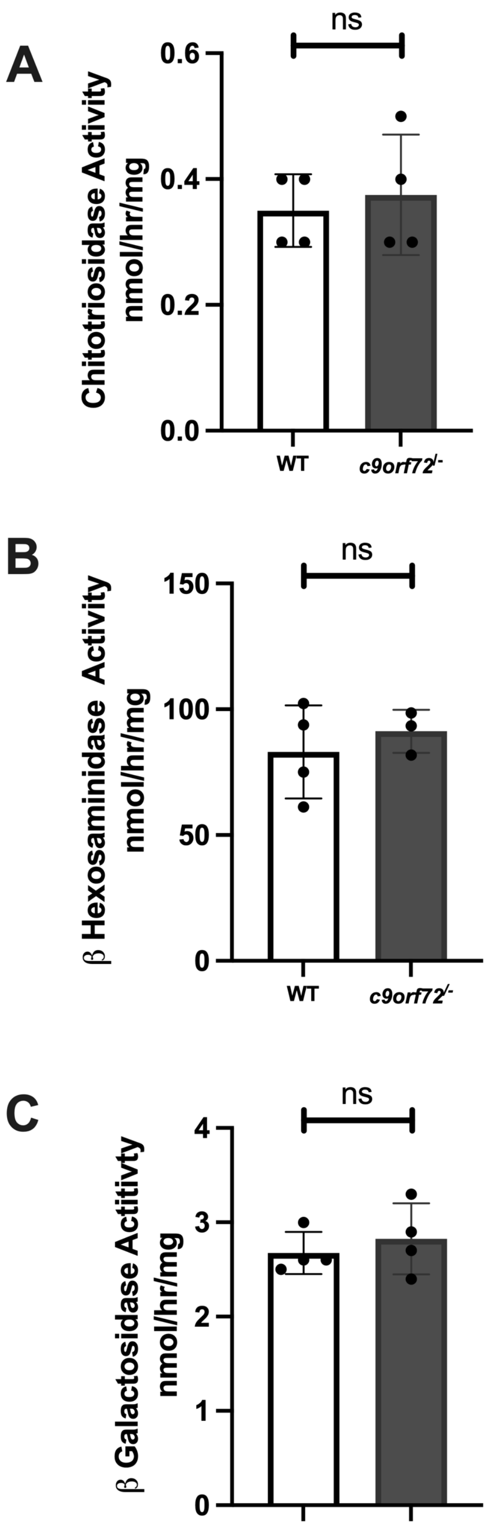


**Supplemental Figure 2. Lysosomal enzyme activities are not significantly altered between *c9orf72*$^{-/-}$ brains and WT controls**. Measurements of different enzyme activities from whole brain homogenates revealed no statistical difference in activity between genotypes. These included Chitotriosidase activity (2A, p=0.6704) a general marker of neuroinflammation and potential ALS biomarker. The lysosomal enzyme enriched in microglia, β Hexosaminidase (2B, p=0.5156) and the lysosomal enzyme β Galactosidase (2C, p= 0.5188). Unpaired, two-tailed and parametric t-test, n=4 for all. These data indicate there is minimal neuroinflammation occurring in the brain.

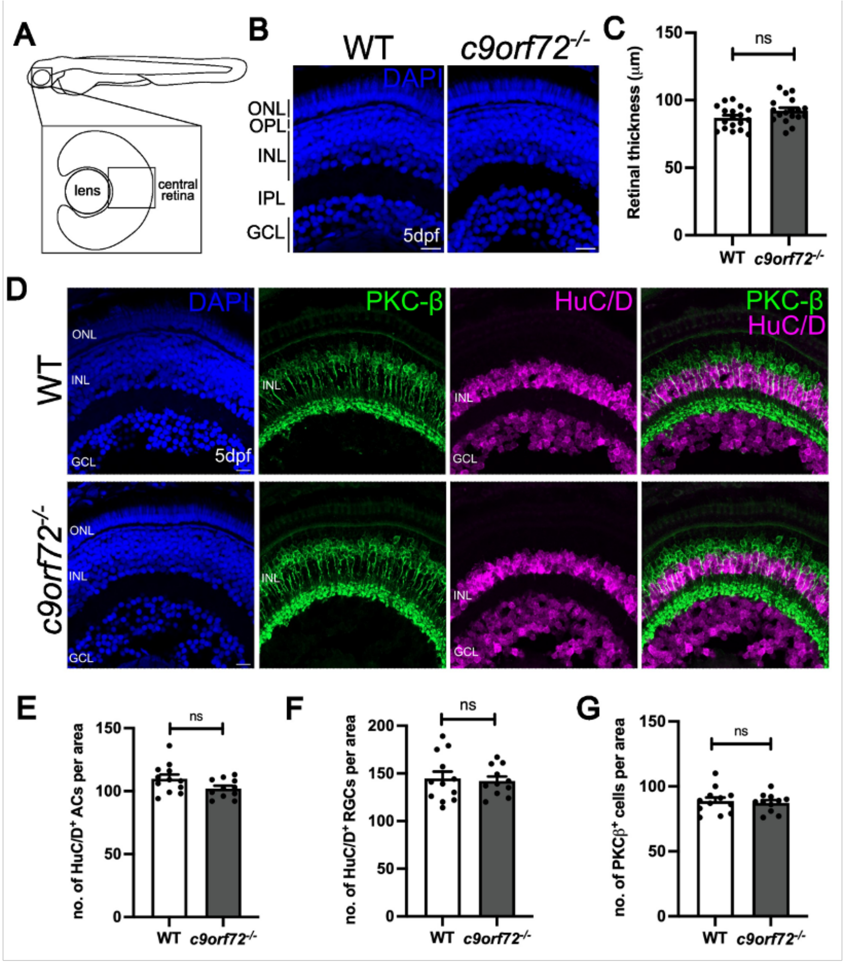


**Supplemental Figure 3. Inner retinal development is normal in *c9orf72*$^{-/-}$ mutants.** A) Schematic diagram of 5-day post fertilisation (dpf) zebrafish retina, highlighting the central retinal region used for analysis. B) DAPI (blue) staining showing the nuclear layers of the WT and *c9orf72*$^{-/-}$ larval retina and overall central retinal thickness. C) Quantification of mean thickness of WT and *c9orf72*$^{-/-}$ retinas; unpaired t-test; p=0.0704. D) Antibody staining for bipolar cell marker, (PKC-β, green), amacrine and retinal ganglion cells (HuC/D, magenta) and nuclei stained with DAPI (blue) in WT and *c9orf72*$^{-/-}$ retinas. D) Quantification of number of HuC/D$^{+}$ amacrine cells in INL per 100□m x 100 x 10□m ROI; unpaired t-test; p=0.0726. E) Quantification of number of HuC/D$^{+}$ retinal ganglion cells in GCL per 100□m x 100 x10□m ROI; unpaired t-test; p= 0.7723. F) Quantification of number of PKC-β$^{+}$ bipolar cells in INL per 100□m x 100 x10□m ROI; unpaired t-test; p= 0.7024; n=11-12 retinas per genotype; Scale bars, 10□m.

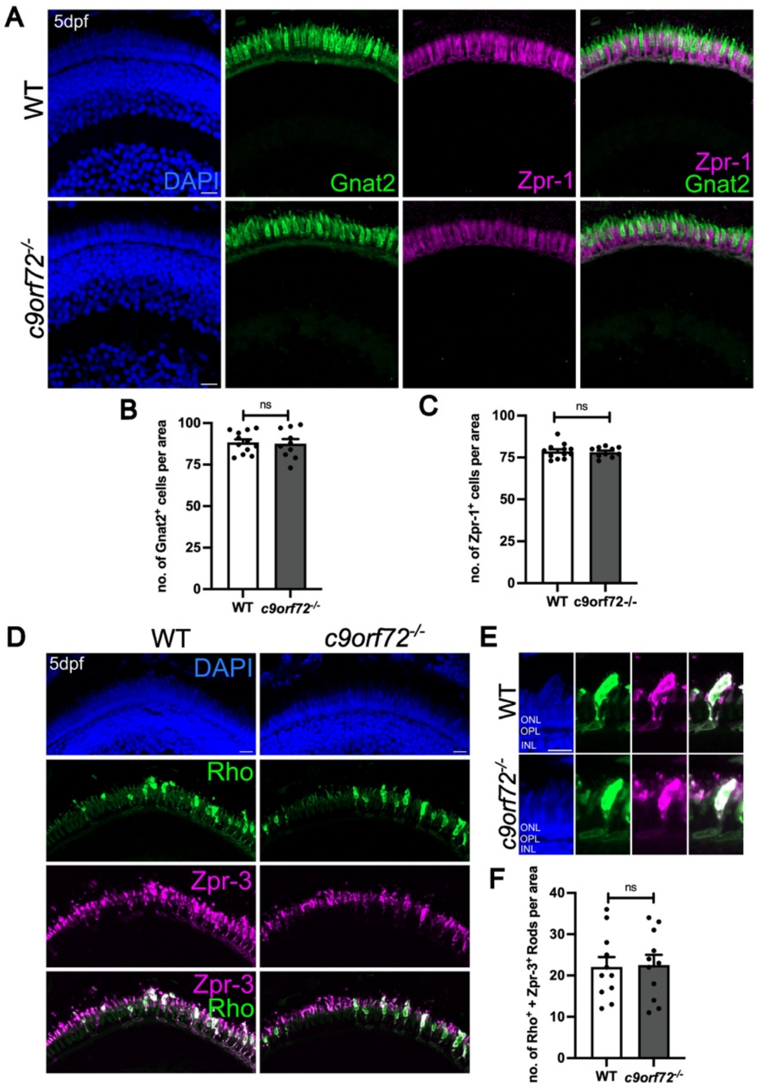


**Supplemental Figure 4. Rod and cone photoreceptors do not exhibit developmental defects in *c9orf72$^{-/-}$ mutants*.** A) Immunostaining for pan-cone marker, Gnat2 (green) and double cone marker, Zpr-1 (magenta) in WT and c9orf72-deficient retinal cryosections at 5dpf. Nuclei labelled with DAPI (blue). B) Quantification of mean number of Gnat2-positive cone photoreceptors in WT and *c9orf72*-deficient retinas;100□m x 100 x 10□m ROI; unpaired t-test, p= 0.8238. C) Quantification of mean number of Zpr-1-positive cone photoreceptors in WT and *c9orf72*-deficient retinas;100□m x 100 x 10□m ROI; unpaired t-test, p= 0.8209. D) Antibody staining for Rhodopsin (Rho; green) and rod/double cone marker Zpr-3 (magenta). E) Close up of individual Rho + Zpr-3-expressing rods in WT and mutants. F) Quantification of mean number of Rho+ Zpr-3$^{+}$ rod photoreceptors in WT and c9orf72-deficient retinas;100□m x 100□m ROI; unpaired t-test, p<0.0001. n=10-12 retinas per genotype. Scale bars, 10□m.

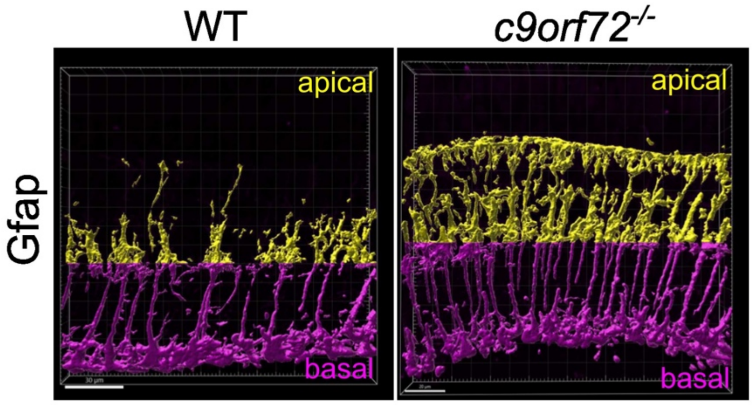


**Supplemental Figure 5. Gfap quantification.** Example of 3D segmentation of Gfap antibody staining in 24-month post fertilisation (mpf) retinal cryosections used for analysis of Gfap distribution along the apicobasal axis. IMARIS was used to quantify the volume of Gfap-staining in the apical (yellow) vs basal (magenta) half of the retina to obtain the ratio of apical:basal Gfap abundance. Left panel: WT; right panel: *c9orf72*.

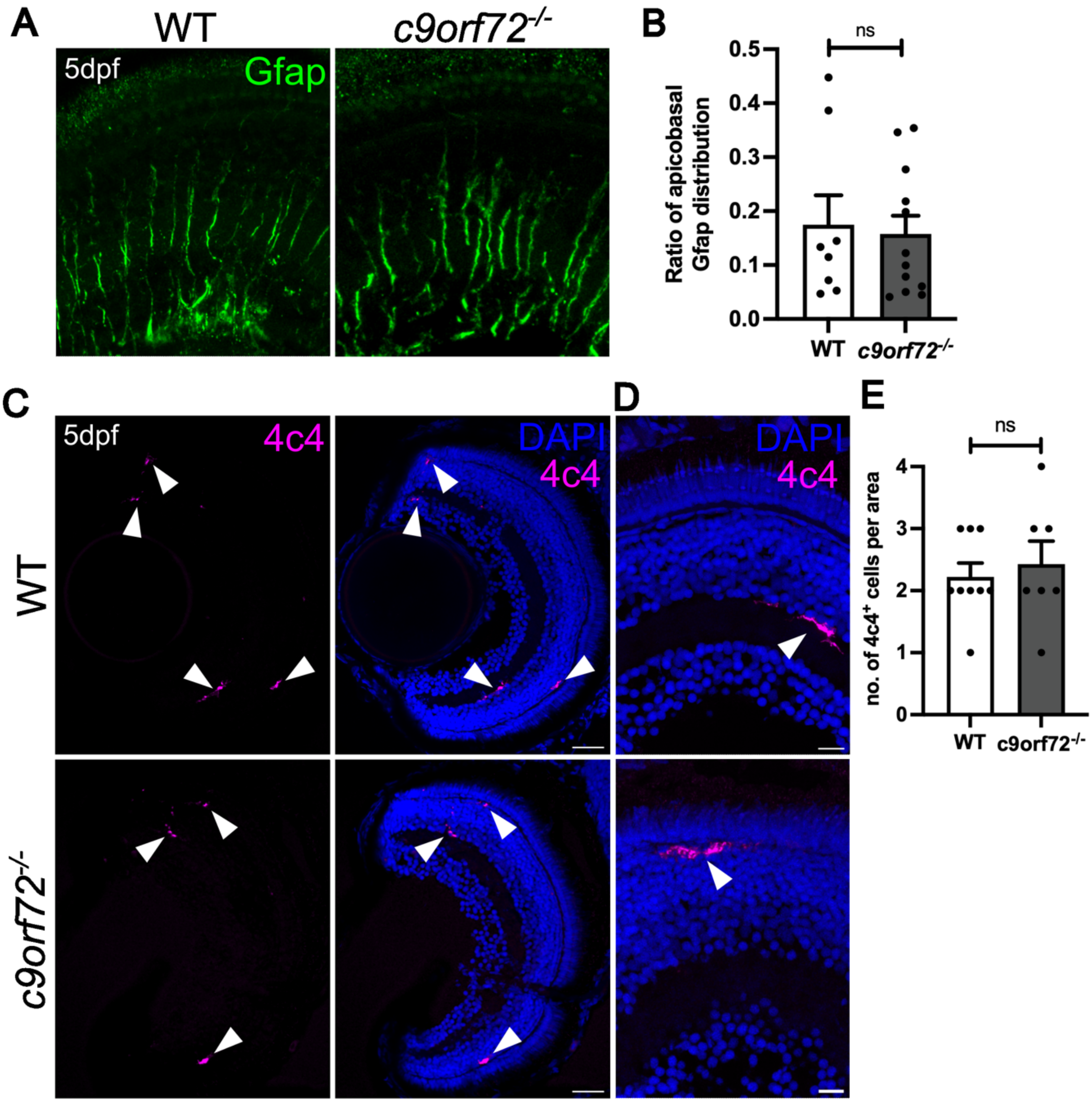


**Supplemental Figure 6. Glial development is unaffected in *c9orf72*$^{-/-}$ deficient retinas.** A) Immunostaining for gliosis marker, Gfap (magenta) in WT and *c9orf72*-deficient retinal cryosections at 5dpf. Nuclei labelled with DAPI (blue). B) Ratio of mean Gfap distribution in apical versus basal regions of the retina; unpaired t-test, p= 0.7791. C) Antibody labelling of retinal microglia with 4c4 (magenta), cell bodies labelled with DAPI (blue). D) Higher magnification image of 4c4 and DAPI staining. E) Quantification of the average number of 4c4$^{+}$ microglia per image; unpaired t-test; p= 0.6226; Scale bars, 10□m.
